# A mechanistic statistical approach to infer invasion characteristics of human-dispersed species with complex life cycle

**DOI:** 10.1101/2024.02.09.578762

**Authors:** Nikunj Goel, Andrew M. Liebhold, Cleo Bertelsmeier, Mevin B. Hooten, Kirill S. Korolev, Timothy H. Keitt

## Abstract

The rising introduction of invasive species through trade networks threatens biodiversity and ecosystem services. Yet, we have a limited understanding of how transportation networks determine patterns of range expansion. This is partly because current analytical models fail to integrate the invader’s life-history dynamics with heterogeneity in human-mediated dispersal patterns. And partly because classical statistical methods often fail to provide reliable estimates of model parameters due to spatial biases in the presence-only records and lack of informative demographic data. To address these gaps, we first formulate an age-structured metapopulation model that uses a probability matrix to emulate human-mediated dispersal patterns. The model reveals that an invader spreads along the shortest network path, such that the inter-patch network distances decrease with increasing traffic volume and reproductive value of hitchhikers. Next, we propose a Bayesian statistical method to estimate model parameters using presence-only data and prior demographic knowledge. To show the utility of the statistical approach, we analyze zebra mussel (*Dreissena polymorpha*) expansion in North America through the commercial shipping network. Our analysis underscores the importance of correcting spatial biases and leveraging priors to answer questions, such as where and when the zebra mussels were introduced and what life-history characteristics make these mollusks successful invaders.

## INTRODUCTION

Biological invasions are a major component of the global environmental crisis. Burgeoning trade and commerce have led to a surge in the establishment and spread of invasive species (Hulme 2009; Meyerson and Mooney 2007), altering the functioning and composition of global ecosystems. Invasions have resulted in trillions of dollars in damage to human health, infrastructure, and production services (Diagne et al. 2021; Turbelin et al. 2023) and are one of the leading drivers of species extinctions (Bellard et al. 2016). Even more alarming is that the pool of invasive species is expected to increase over the next century as new trade routes emerge (Seebens et al. 2018). Therefore, a major theoretical and practical challenge in invasion science is to understand how human transportation patterns interact with the life-history dynamics to determine the spread of an invasive species (Lewis et al. 2016).

Early efforts to model invasions were based on integrodifference equations (IDE) (Kot et al. 1996; Lutscher 2019) that used fat-tailed redistribution kernels to emulate long-distance dispersal by human transportation (Hallatschek and Fisher 2014). The IDE models are conceptually attractive because they can be combined with structured demographic models (Neubert and Caswell 2000) and permit analytical solutions (Bateman et al. 2015). This allows researchers to understand invasion patterns intuitively and provides valuable insights into managing invaders with complex life histories. For example, a Gaussian IDE model predicts a radially expanding invasion front (Kot et al. 1996), such that the species’ arrival time is a linear function of distance from the origin (Skellam 1951). Moreover, managers can conduct sensitivity analyses to identify which life-history parameters should be targeted to slow or stop invasion (Neubert and Caswell 2000).

Despite the conceptual simplicity of IDE models, they have two major limitations. IDE models assume that species dispersal is directionally random (i.e., isotropic) (Thompson et al. 2021), and the shape of the dispersal kernel does not change from location to location (i.e., spatially invariant) (Hallatschek and Fisher 2014). However, human-mediated dispersal patterns are highly heterogeneous owing to the complex topology of transportation networks (Banks et al. 2015). Transportation networks are characterized by a local cluster of densely connected neighbors and a few long-distance connections between clusters. This feature creates a small world structure (Hu and Zhu 2009), leading to anisotropy and spatial invariance in dispersal patterns (Wolf et al. 2019). Consequently, the patterns of invasions appear spatially incoherent, and, as such, distance from the origin is a poor predictor of species arrival time (Horvitz et al. 2017).

These limitations of IDE models are well recognized (Hallatschek and Fisher 2014; Thompson et al. 2021), and several mathematical approaches have been developed to address them (Hui and Richardson 2017). Among these approaches, metapopulation models are gaining popularity, in part due to their success in describing the spread of infectious diseases through human transportation networks (Brockmann and Helbing 2013; Colizza et al. 2006; Gautreau et al. 2008; Iannelli et al. 2017; Rvachev and Longini Jr 1985). Broadly, metapopulation models consist of discrete patches (such as a lake or a county) within which the population changes according to a demographic model and edges that connect these patches to capture heterogeneous dispersal patterns of invaders. Because of this flexibility, metapopulation models have been used to study the expansion of invasive species through a wide range of human transportation mechanisms (Carrasco et al. 2010; Chapman et al. 2016; Ferrari et al. 2014; Floerl et al. 2009; Seebens et al. 2019).

However, these implementations of metapopulation models are conceptually impenetrable because researchers often err on the side of realism by creating parameter-rich computational models that incorporate minute species and transportation-specific details. As a result, it becomes challenging to decipher what aspects of species biology and transportation networks determine the broad-scale patterns of invasions and what are the general underlying ecological principles that characterize these patterns (Brockmann and Helbing 2013; May 2004).

Building simple conceptual models of human-mediated invasions is only one part of the challenge. To make these models useful, we also need statistical methods to estimate model parameters to answer questions such as—where and when the species was introduced and what makes a given species a successful invader. Answering such questions is critical for formulating management actions to contain and eradicate invaders. However, estimating model parameters is challenging because species distribution data are often fraught with errors and biases (Isaac et al. 2014; Johnston et al. 2023). For instance, most commonly available species occurrence records are gathered by opportunistic sampling rather than structured surveys that report both the presence and absence of species. Such types of unstructured data, referred to as presence-only data, typically suffer from irregular spatial sampling, making it hard to narrow down the location of species introduction. Even when sampling is structured, a species may remain undetected during the establishment phase, making it challenging to infer the time of species introduction (Crooks 2005).

Another potential issue in estimating parameters may arise if multiple combinations of two or more parameters produce the same ecological outcome. For example, two species with different life-history strategies can generate the same spread patterns (Bateman et al. 2015). This issue of non-identifiability of parameters makes it statistically challenging to infer the life-history tactics of an invader, which, in turn, can limit our ability to determine which life-history parameters play an outsized role in the species spread and, therefore, should be the target of management efforts (Crouse et al. 1987).

Recent theoretical and computational advances in applied Bayesian statistics provide avenues to overcome these hurdles (Gelman et al. 1995; Hobbs and Hooten 2015; Hooten and Hefley 2019; McElreath 2020). Bayesian models learn parameters using a data model and a parameter model. The data model (or the likelihood) is a probabilistic generative model of observed data. For invasions, this generative model is a multi-level process in which the occurrence data are linked to latent parameters, such as invaders’ abundance, via an observation model that describes how the data were collected. These abundances are then, in turn, linked to demographic parameters via a mechanistic model of invasion (Hooten et al. 2007; Wikle 2003). This hierarchy offers a key advantage—the information from data to demographic parameters flows through an observation model where one can explicitly incorporate the spatial biases and detection errors in the sampling procedure. This may allow one to obtain a reliable estimate of parameters, such as where and when the species was introduced in the alien territory, even when sampling is spatially unstructured.

In contrast to the data model, the parameter model (*i*.*e*., prior) characterizes our uncertainty of the parameters before the data are collected. Intuitively, a parameter model is an expression of our prior scientific knowledge. For example, population biology dictates that a successful invader will have a positive growth rate. Therefore, we can constrain the growth rate to take only positive values *a priori*. Similarly, a biologist may have additional information about the invader, such as its life-history parameters, based on previous studies on the invader in its native range or a closely related species. By including this information via a parameter model, Bayesian models can disentangle otherwise non-identifiable parameters (Neath and Samaniego 1997).

In this study, we aim to fill the aforementioned theoretical and statistical gaps to understand the spread of invasive species by human movement. We first formulate an analytically tractable age-structured metapopulation model to gain a conceptual understanding of how complexity in human transportation networks determines the spatiotemporal patterns of species spread. We ask how properties of the transportation network and species life-history tactics determine the invasion path and arrival time. Next, we propose a hierarchical Bayesian model to estimate the parameters of the metapopulation model using presence-only data and prior scientific knowledge. We tailor our mathematical and statistical models to analyze the expansion of zebra mussels (*Dreissena polymorpha*) in the inland waterways of North America via the commercial shipping network. We identify the time and place of zebra mussel introduction, the invasion route, and the life-history parameters that are most critical for its spread. Despite our focus on zebra mussels, the conceptual insights offered by the metapopulation model and the statistical guidance to analyze empirical data are general and broadly applicable.

### Zebra Mussels

Zebra mussels are freshwater bivalves native to Eastern Europe and were likely introduced in the Great Lakes by the release of contaminated ballast water (Nalepa and Schloesser 2013). Post introduction, zebra mussels have dramatically altered North America’s water ecology and local economy. These mollusks are filter feeders that primarily consume planktonic algae and microzooplankton, thereby changing the energy flow and structure of food webs in water bodies where they reside. They are notorious biofouling organisms (Ludyanskiy et al. 1993)—their active byssal apparatus allows them to attach to hard substrates at very high densities that can impede the function of human infrastructure, resulting in an annual loss of half a billion dollars (Warziniack et al. 2021). Due to these negative impacts on biodiversity and the economy, zebra mussels have earned a spot in the International Union for Conservation of Nature’s list of 100 of the world’s worst invaders (Lowe et al. 2004). Therefore, understanding the expansion characteristics of zebra mussels can help us slow or stop its spread and may provide insights into managing other emerging invasions.

However, due to detection errors and biased sampling of zebra mussels, tracing the invasion pathway and identifying the time and place of their introduction is challenging. They were first detected during a large-scale survey in 1988 in Lake St. Clair (Hebert et al. 1989). However, the rediscovery of a 1986 letter exchange between Pembina Explorations Ltd. and the Ontario Ministry of Natural Resources suggests that a colony of mussels was present in Lake Erie near Kingsville, Ontario, Canada (Carlton 2008). In subsequent years, zebra mussel detections spiked at distant locations across the Great Lakes (GBIF.org 2022; USGS 2022) (Fig. S1). By 2010, they were detected and established in most of the inland commercial waterways from the Great Lakes to the Gulf of Mexico, implicating hitchhiking by hull fouling of barges and tugs as putatively the primary mechanism of continental-scale dispersal (Johnson and Padilla 1996). These established populations in commercial waterways are, in turn, fueling over-land range expansion to isolated water bodies through recreational boating (Johnson and Carlton 1996).

## RESULTS

### Formulation and Results of the Metapopulation Model

To model the continental-scale range expansion of zebra mussels in North America, we consider an environmentally homogeneous metapopulation with *K* patches (or locations) connected by the commercial shipping network. For zebra mussels, these locations can correspond to a port or a collection of ports in some fixed area. At each location, the zebra mussel population is structured by age, such that the population dynamics are governed by the following matrix difference equation (Hunter and Caswell 2005):

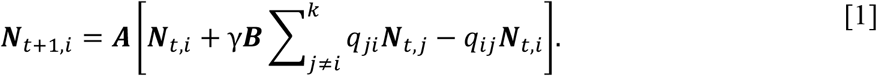

***N***_*t,i*_ is a vector of the population in different age classes at location *i* and time *t, γ* is the total number of voyages that ships take in one-time step, ***B*** is a diagonal matrix with elements corresponding to age-dependent hitchhiking rate, and ***A*** is the Leslie matrix that captures age dependent life-history schedules of zebra mussels such as reproduction, survival, and mortality (Leslie 1945). When the invader population is structured by ontogenetic stages, we replace the Leslie matrix with the Lefkovitch matrix to capture stage-dependent life-history events (Lefkovitch 1965). Dispersal of propagules is weighted by *q*_*ij*_, where *q*_*ij*_ is the fraction of voyages from patch *i* to patch *j*, such that **∑**_*i*_ **∑**_*j ≠ i*_ *q*_*ij*_ = 1.

Using port call data from MarineTraffic, we estimate weights, *q*_*ij*_, at 50 km resolution and simulate the hypothetical spread of zebra mussels originating near Kingsville (Ontario, Canada) (brown point in Fig. 1C), where mussels were first detected in late 1986 (Carlton 2008). We make two key observations. First, time to species arrival is correlated with river distance from Kingsville, but with substantial variation from the best-fit line (R^2^ = 0.89; Fig. 1A). Second, the spatiotemporal patterns of species colonization are incoherent (Fig. 1C), exhibiting only a weak trend toward the equator. These observations sharply contrast the general predictions of the Gaussian IDE model (Kot et al. 1996), which produces a radially expanding wavefront with a linear relationship between arrival time and distance from the point of introduction (Skellam 1951).

**Figure 1:**
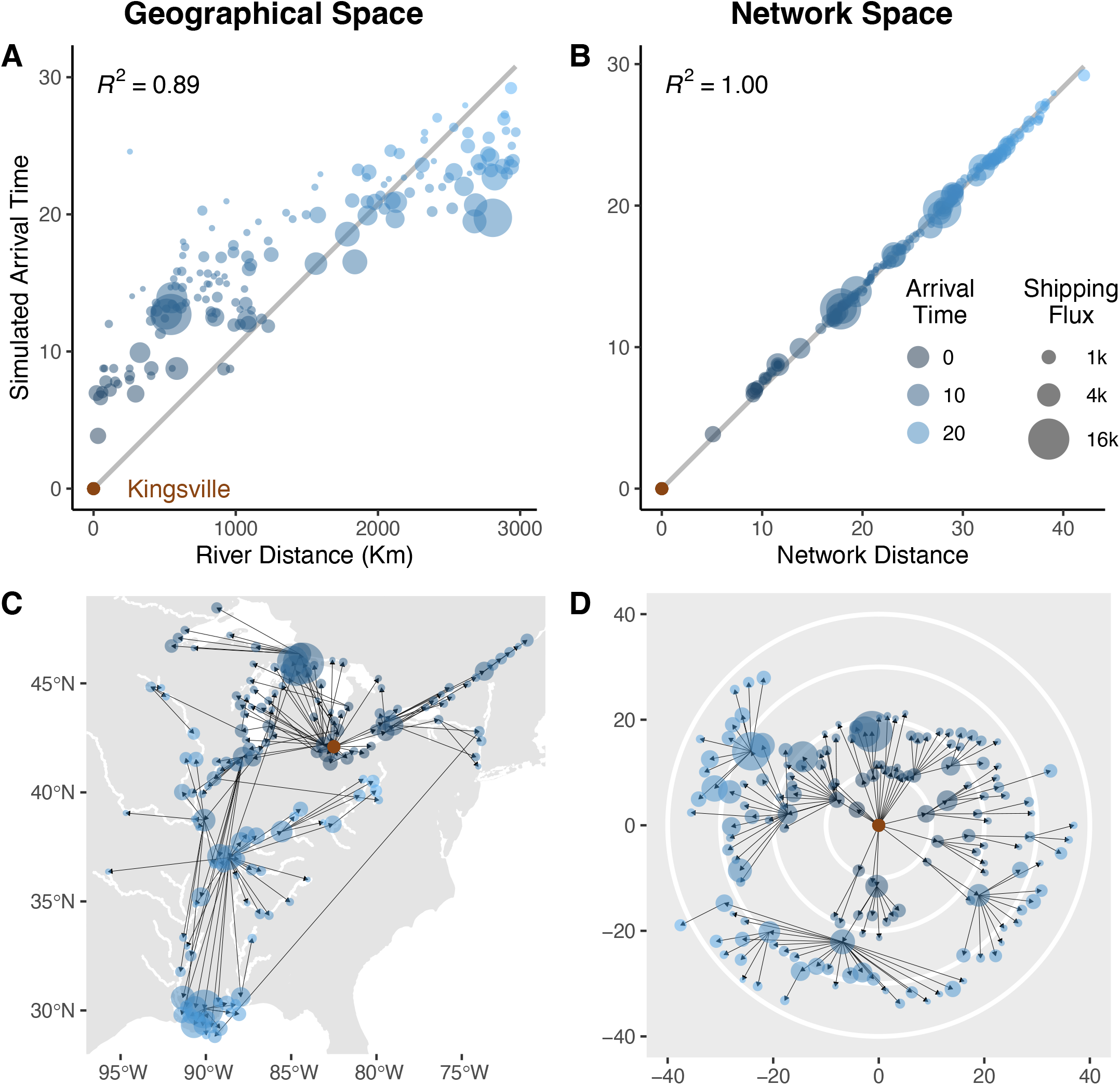
A deterministic simulation of the spread of zebra mussels originating from Kingsville, Ontario (brown point) on the geographical and network space. On the geographical space, (**A**) the river distance from Kingsville is a relatively weak predictor of the time of zebra mussel arrival and (**C**) the pattern of expansion is spatially incoherent. In contrast, on the network space, (**B**) the shortest network distance from the origin (Eq. [9]) shows an almost perfect linear association with the species arrival time (Eq. [10]). (**D**) Visualizing species spread on the network space yields radially expanding wavefront—reminiscent of IDE models (Kot et al. 1996).

The discrepancy in the arrival statistics of the metapopulation model from Gaussian IDE models shows that the conventional approach to model human-mediated invasion may be limited. We reformulate the invasion dynamics to highlight the hidden underlying structure we can exploit to understand the spread of invasive species via human transportation networks (Brockmann and Helbing 2013; Iannelli et al. 2017). To do that, we assume that for human-dispersed species, the dispersal inputs to a population contribute much less than local reproduction. This allows us to approximate ***N***_*t,i*_ as a scalar multiple of the stable-age distribution (***W***; proportion of individuals in each age class), ***N***_*t,i*_ ≈ *x*_*t,i*_***W***, where

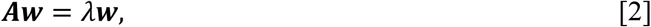

and *λ* is the species growth rate (Caswell 2001). Intuitively, this approximation allows us to project equation [1] onto ***W***, which simplifies the matrix difference equation to a scalar difference equation:

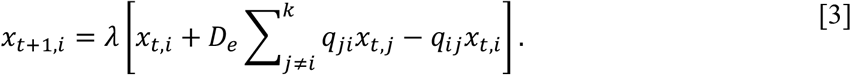

In Eq. [3],

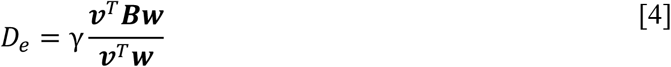

is effective dispersal, and ***v***^*T*^ is the reproductive value (total number of offspring born to an individual from their current age onwards), which is given by

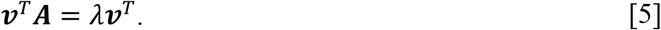

We use the superscript *T* to denote the transpose of a vector or a matrix (see supporting information).

Note that the scalar approximation effectively partitions the metapopulation model (Eq. [1]) into two hierarchical levels—spatial and life-history dynamics. These levels interact via population-level parameters *λ* and *D*_*e*_, where *λ* is the dominant eigenvalue of the Leslie matrix (Eqs. [2] and [5]) and *D*_*e*_ is the mean age-specific propagule pressure weighted by the reproductive value (Eq. [4]). This link between population level (*D*_*e*_ and *λ*) and life-history parameters (***A*** and ***B***) highlights the importance of incorporating age or stage structure in invasion models (Neubert and Caswell 2000; van den Bosch et al. 1992; Van den Bosch et al. 1990) because it may sometimes produce counterintuitive predictions. For example, in an extreme case where the invader disperses post-reproductive senescence, the species will fail to spread because effective dispersal (*D*_*e*_) is zero even though both propagule pressure and growth rate are positive.

To simplify the model further, we define 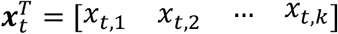, which allows us to express the joint population change in all locations as

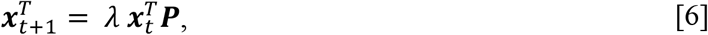

where ***P*** is a probability matrix. The non-diagonal elements of the matrix, *P*_*ij*_ = *D*_*e*_*q*_*ij*_, correspond to the proportion of mussels dispersing from patch *i* to *j* in one time step. Note that the structure of the metapopulation model in Eq. [6] bears a remarkable resemblance with IDE models (Kot et al. 1996). In both models, the population at the next time step is a product of population growth rate and dispersal. In IDE models, dispersal is captured using a redistribution kernel, which integrates to unity. Similarly, in the metapopulation model, we capture dispersal using a probability matrix ***P*** (hereafter referred to as the redistribution matrix) in which each row sums to unity. But, unlike redistribution kernels, the redistribution matrix can capture spatial invariance and anisotropy in dispersal patterns.

Next, assuming 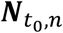 mussels are introduced at location *n* at time *t*_0_, the population after *t* time steps is given by

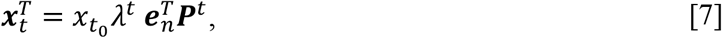

where 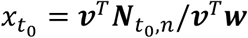 is the effective initial population, and 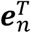 is a row vector with all zero elements except at index *n*, where it takes value one. The population change is a product of exponential growth, 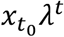 and a *t*-step transition probability, 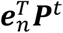 that encapsulates the dispersal trajectory of zebra mussels by the inland shipping network (Kulkarni 2016). Intuitively, 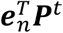 is the probability of finding a zebra mussel at a given location after *t* time steps via any path originating from *n*.

A natural question is whether multiple paths contribute to zebra mussel arrival because they might be equally likely or if a single path has the dominant contribution to the expansion process? And, if the latter were the case, what would determine this path? In principle, the invader can arrive at a location by infinite paths. However, not all paths are equally probable. Human transport networks are highly heterogeneous, so there is usually a single fastest path to the destination (Brockmann and Helbing 2013; Gautreau et al. 2008).

To understand what determines this path, we define network distance from location *i* to *j* as

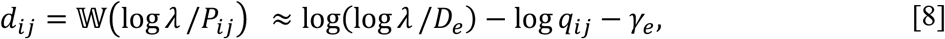

where 𝕎 is the Lambert W function, and *γ*_*e*_ is the Euler-Mascheroni constant (Gautreau et al. 2008). This definition of distance emerges naturally from the theory of discrete stochastic processes (Kulkarni 2016) and provides an intuitive way to visualize and understand human-mediated invasions. For one, this notion of distance suggests that, on the network space, two locations are close if they exchange a high traffic volume and the hitchhikers have high reproductive value (*i*.*e*., high *D*_*e*_).

Now consider a set of all paths from *n* (origin) to *m* (destination)— Γ _*nm*_. We postulate that the zebra mussel will arrive at *m* via the shortest network path 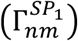, which maximizes the product of dispersal probabilities along the path (Bagnara et al. 2022; Brockmann and Helbing 2013; Gautreau et al. 2008; Iannelli et al. 2017). As such, the total network distance traveled is given by

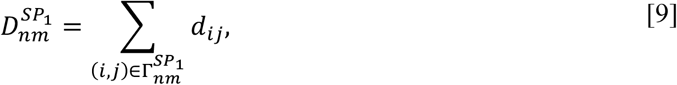

and the time of the species’ arrival is

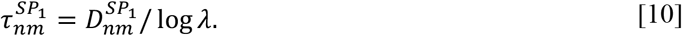

Although mussels can arrive by alternate paths, their contribution diminishes exponentially with increasing network distance such that the shortest path is the major contributor to the most probable distance,

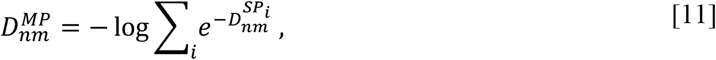

where 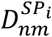 is the network distance along the *i*^th^ shortest path from *n* to *m* (Iannelli et al. 2017).

Indeed, simulations confirm a strong linear relationship between the shortest network distance from the origin and arrival time (*R*^2^ = 1.00; Fig. 1B), with a mean error of about 2% when we consider all possible invasion paths (Fig. S2A). The linear relation also holds even when we consider stochastic dispersal, reproduction, and survival (Fig. S2B). These findings explains why the physical distances are a relatively weak predictor of species arrival time (Padilla et al. 1996) (Fig. 1A). And, interestingly, visualizing species spread on the network space yields a radially expanding wavefront (Fig. 1D)— reminiscent of IDE models—even though on the geographical space, the spread patterns are incoherent (Fig. 1C) (Brockmann and Helbing 2013).

### Formulation of the Hierarchical Bayesian Model

To estimate the parameters of the metapopulation model, we specify a hierarchical Bayesian model with three levels—(*a*) an *observer model* that describes how variation in ’true’ abundance translates to invader presence data; (*b*) a *spatial dynamics model* that describes spatiotemporal variation in ’true’ abundances of the invader; (*c*) a *life-history model* to describe survivorship, fecundity, and dispersal schedules of the species. These sub-models form a hierarchy in that the output of the life-history model is the input for the spatial dynamics model, whose output is the input of the observer model (Pagel and Schurr 2012).

In a Bayesian framework, this hierarchy is implemented by partitioning the likelihood into the product of probabilities corresponding to each sub-model by conditioning (Berliner 1996). Then, using the Bayes rule, the posterior distribution of parameters is expressed as the product of the likelihood (informed by data) and the prior (informed by beliefs based on expert opinion or scientific experiments conducted by others). This posterior is a high-dimensional joint probability distribution of parameters, which is used to infer parameter expectations and uncertainties (Gelman et al. 1995). We follow the recommended Bayesian workflow guidelines to specify the model, evaluate the output, and make inferences (Gelman et al. 2020). We outline the main components of the three sub-models in Figure 2 and summarize how these sub-models are linked hierarchically in the section below. For the full statistical model, including implementation details statements, relationship between parameters, and priors, see Table S1.

**Figure 2:**
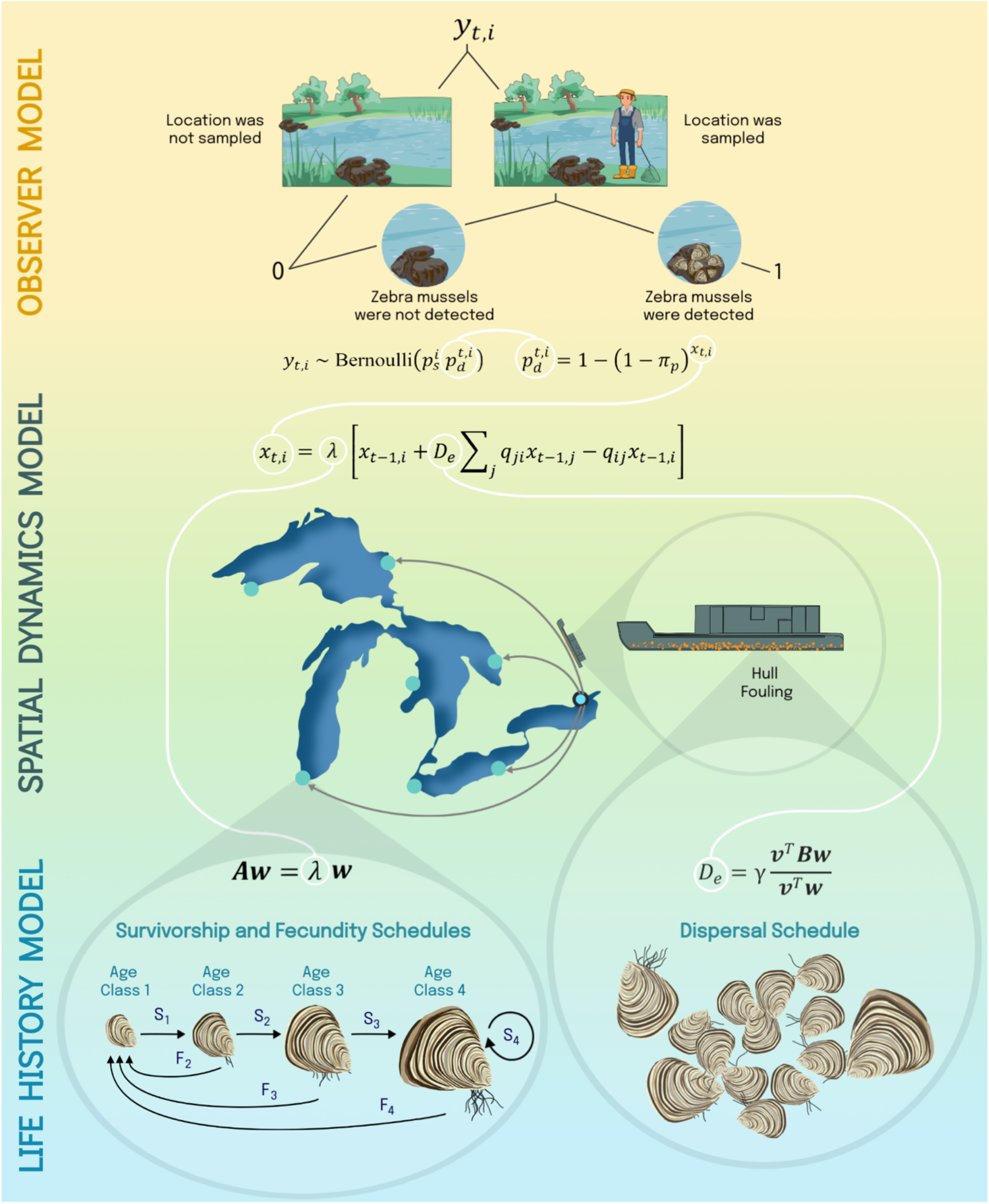
A conceptual diagram describing the hierarchical structure of the Bayesian model. The observer model (*top*) links the presence data, *y*_*t,i*_, with zebra mussel abundance, *x*_*t,i*_ (Eqs. [12] and [15]). The spatial dynamics model (*middle*) describes population change due to local growth and dispersal by the shipping network (Eq. [16]). Finally, the life-history model (*bottom*) describes how zebra mussel survival, fecundity, and dispersal schedules (***A*** and ***B***) determine the population level parameters—growth rate, *λ*, and effective dispersal, *D*_*e*_ (Eqs. [2], and [4]). These sub-models form a hierarchy in the sense that the output of the lower level is the input of the level above (Pagel and Schurr 2012). The illustration was made by Upasana Sarraju.

#### Observer model

To characterize the presence-only data for zebra mussels (GBIF.org 2022; USGS 2022), we define an indicator data variable *y*_*t,i*_ that takes value one if mussels are reported within a 25 km radius of location *i* at time *t* and zero otherwise. When *y*_*t,i*_ = 1, the location is sampled, and mussels are detected. However, when *y*_*t,i*_ = 0, it can imply one of two things—either a location was not sampled or, when sampled, the mussels were not detected. Implicitly, this data model neglects false positives, which may be reasonable as these datasets are subject to quality controls before cataloging. We model this data-generative process as a mixture with two components. Location *i* is sampled with probability 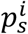 and, given that the location is sampled, mussels are detected with probability 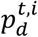 (see observer model in Fig. 2). Cumulatively, this yields

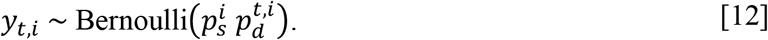

We assume sampling probability is constant over time but can vary with location, such that nearby locations have similar sampling probabilities. To account for this spatial information, we model the joint distribution of sampling probabilities across locations as a multivariate normal distribution,

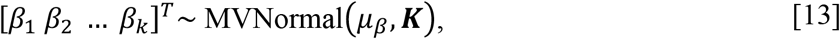

where *logit*^−1^(*β*_*i*_) is the sampling probability at location *i, logit*^−1^(*μ*_*β*_)is the mean sampling probability on the logit scale, and *K*_*ij*_ is the covariance between location *i* and *j*. We use an RBF kernel,

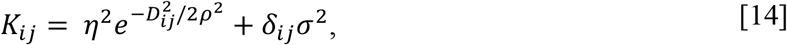

to model covariance, *D*_*ij*_ is the geographical distance between locations *i* and *j*, and *δ*_*ij*_ is the Kronecker delta function. The hyperparameter *ρ* is the length scales of correlation, *η* is the maximum covariance between two locations, and *σ* provides extra covariance when *i* = *j*. Modeling spatial correlation in sampling probabilities using a multivariate normal distribution allows partial pooling, thus providing a more robust estimate of sampling probabilities for locations where data are insufficient. Assuming *π*_*p*_ is the permussel detection probability, the detection probability of the species at location *i* and time *t* is

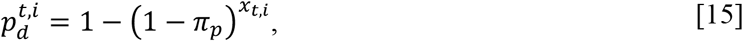

where *x*_*t,i*_ is the local abundance of mussels.

#### Spatial dynamics model

To model changes in abundance of mussels (via all routes), we use the spatial dynamics model in Eq. [6] (see spatial dynamics model in Fig. 2), with a slight modification—at high density, the population saturates to the carrying capacity. We achieve this by reformulating the spatial dynamics model as

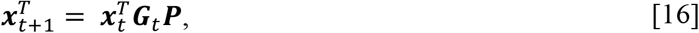

where 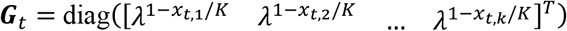 is the Ricker growth matrix (Ricker 1954) with carrying capacity *K*. Although density dependence does not affect arrival statistics because of exponential growth at the expanding front, it helps avoid numerical pathologies during inference. Recall, *P*_*i*≠*j*_ = *D*_*e*_*P*_*i*_*q*_*ij*_, where *D*_*e*_ = *γ****v***^*T*^***BW***/***v***^*T*^***W***. We assume 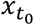 mussels are introduced at *n* such that 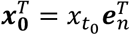 Next, by setting the effective initial population to one (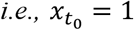) we can estimate the earliest time of introduction. Unfortunately, the nature of the observation and spatial dynamics models do not allow us to infer *t*_0_. At carrying capacity, the detection probability is approximately 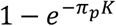 Therefore, even though the product of *π*_*p*_ and *K* is identifiable, these parameters are individually non-identifiable, which implies that we can only estimate local mussel abundances relative to carrying capacity. In other words, we cannot estimate 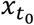 and, consequently, cannot identify *t*_0_. We can remedy this situation through a parameter model by constraining carrying capacity. Studies estimate that the carrying capacity of quagga mussels (a species similar to zebra mussel in its biological characteristics) varies between 10 and 100 thousand individuals per m^2^ (Cross et al. 2011). Using quagga mussel carrying capacity as a prior for zebra mussel carrying capacity, we can calculate the time, 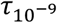 when the population at origin is 10^−9^*K* or 1-10 individuals per km^2^. Assuming 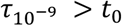, we use 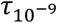 as a proxy for the time of zebra mussel introduction.

#### Life-history model

We model local patch dynamics using a Leslie matrix ***A*** with four age classes of width one year:

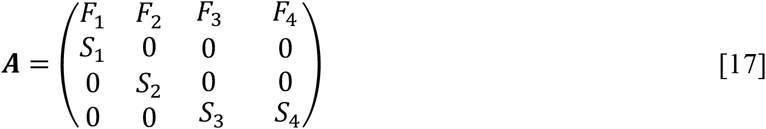

(see the life-history model in Fig. 2). Similar to carrying capacity, the presence records cannot constrain the elements of the Leslie matrix because multiple combinations of life-history parameters yield the same *D*_*e*_ and *λ*. We circumvent this limitation by assigning informative priors on age-specific survivorship probability (*S*_*i*_) and fecundity (*F*_*i*_) based on estimates from literature (Casagrandi et al. 2007) (Table S1). We also constrain the estimates of age-specific hitchhiking rate (***B***) using published data on the shell length of randomly sampled zebra mussels on a commercial vessel (Keevin et al. 1992). Based on the allometric relationship between zebra mussels’ age and shell length, the data suggest that attached mussels were older than one year. Therefore, we assume that the hitchhiking rate is very low for age class 1 (*B*_11_ ≈ 0) and remains constant after (*B*_22_ = *B*_33_ = *B*_44_ = *B*). We also consider an alternate hypothesis of constant hitchhiking rate for all age classes (Fig. S3).

### Results of the Hierarchical Bayesian Model

We implement the above statistical model in the Stan language (Carpenter et al. 2017). Internally, Stan samples from a probability distribution using no-U-turn sampler (Hoffman and Gelman 2014), which is an efficient implementation of the Hamiltonian Monte Carlo MCMC algorithm (Neal 2011) (see supporting information for Stan and R code).

#### Model testing

The choice of prior can play a notable role in Bayesian models (Gelman et al. 2020). On the one hand, strong priors can resolve non-identifiable parameters such as the elements of the hitchhiking and Leslie matrix. On the other hand, vague priors on transformed parameters can lead to unrealistic expectations on the outcome scale. For example, vague hyperpriors of the covariance function (Eq. [13]) can lead to a highly informative U-shape prior on the probability scale, thereby biasing estimates of sampling probabilities. For these reasons, we evaluate and refine our priors by simulating invasion outcomes using parameters drawn from the prior distribution. Our choice of prior yields biologically realistic invasion outcomes and parameter values.

Next, to test if the statistical model works as intended, we generated synthetic data according to the generative model described above (Eqs. [12-17]) and then fit the statistical model to the synthetic data. Diagnostics reveal that the MCMC chains converge and yield reliable estimates of means and interquartile ranges. For most parameters, the actual value lies within the 95% credible interval. Posterior predictive checks confirm that the predicted time of first detection (Fig. 3A) and the number of zebra mussel detections at a location (Fig. 3C) are consistent with synthetic data. Importantly, we can reasonably predict the time (using 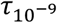 as a proxy; Fig. 4A) and the location of the introduction (Fig. 4C; actual values are marked with an asterisk).

**Figure 3:**
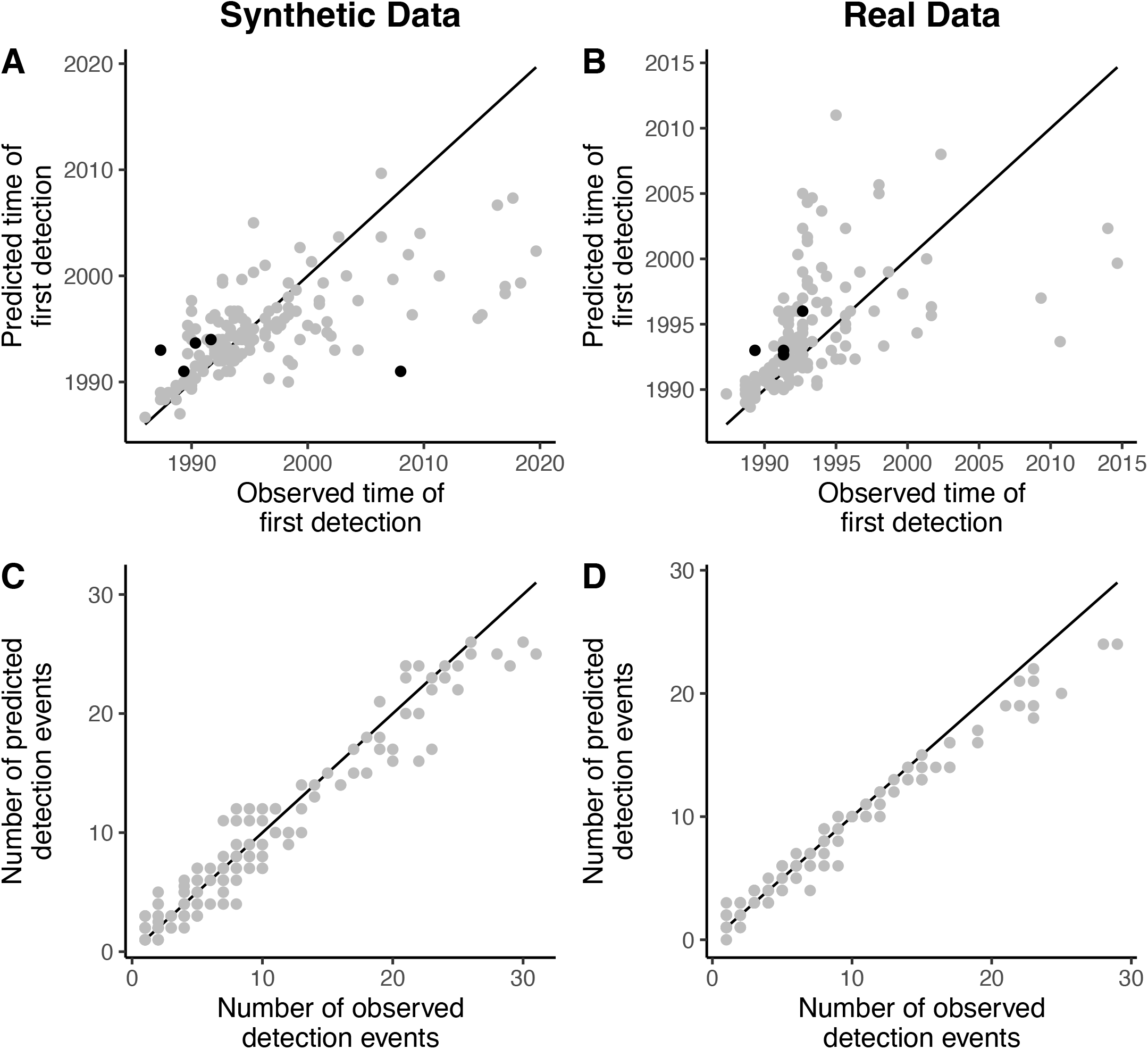
Posterior predictive checks for synthetic data (left column) and real data (right column) suggest that the predicted time of first detection (**A** and **B**) and the number of detections of zebra mussels (**C** and **D**) at a particular location are consistent with the observed time of first detection and the number of confirmed detections, respectively. In both plots, the points represent the median value of the estimate. For grey (black) points, the observed data lies inside (outside) the 95% credible interval. The black line represents the perfect match between the data and the statistical estimate.

**Figure 4:**
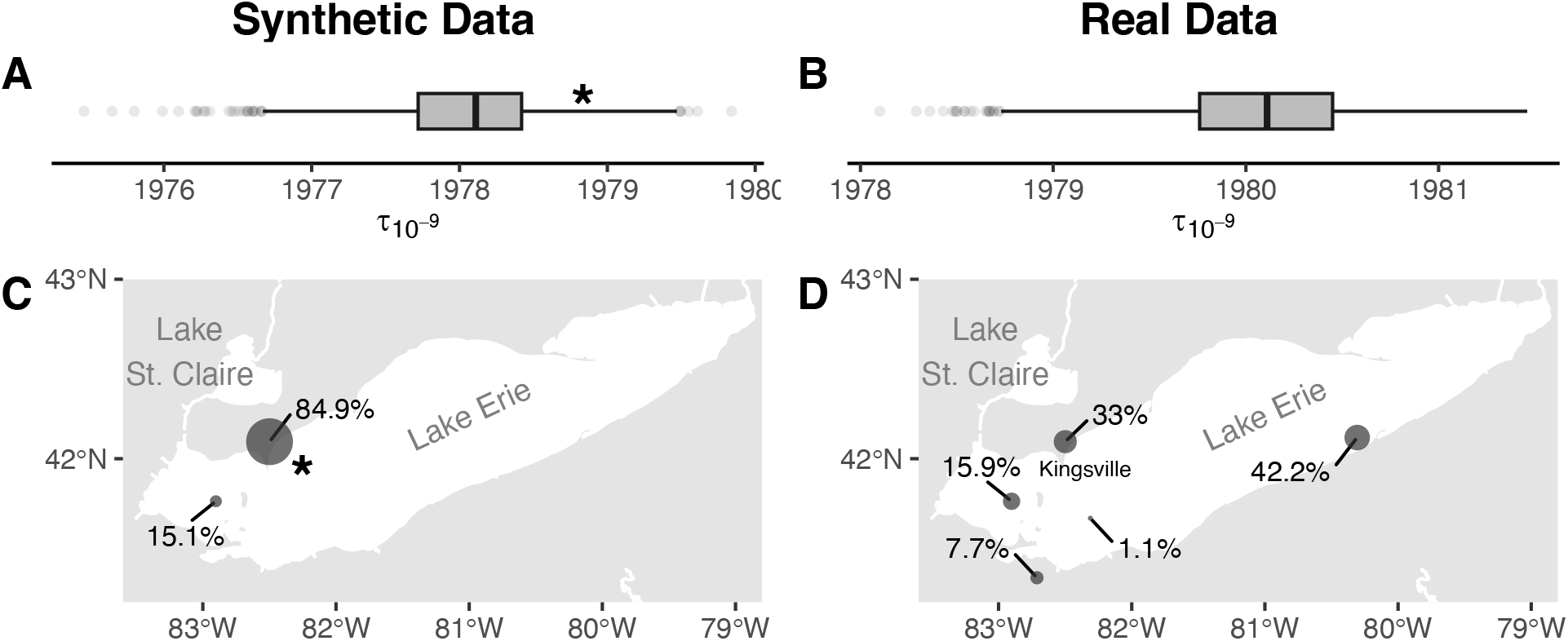
The plots show the posterior distribution of time (using 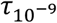 as proxy) and place of zebra mussel introduction for synthetic (left column) and real data (right column). Fitting the Bayesian model to synthetic data generated from known parameter values recovers the 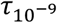 (**A**) and location of the introduction (**C**) (true parameter values are marked with an asterisk). Analyzing the real data suggests that zebra mussels were likely introduced before 1981 in Lake Erie (**B**)—at least five years before the first confirmed detection in late 1986 (Carlton 2008). However, within Lake Erie, the statistical model identifies multiple locations where the mussels could have been introduced (**D**). Interestingly, there is only one in three chance that mussels were introduced in Kingsville, where they were first detected in 1986.

### Data analysis

We fit the statistical model to zebra mussel occurrence data recorded from Jan 1980 to Dec 2019 (GBIF.org 2022; USGS 2022). Diagnostics reveal that the MCMC chains converge and yield reliable estimates of parameter means and interquartile ranges. To evaluate the model fit, we performed posterior predictive checks. We found that the predicted time of first detection and the number of zebra mussel detections at a location are consistent with real data (Figs. 3B and 3D).

The model fit reveals intriguing features about the initial phases of zebra mussel expansion. The bulk of the posterior distribution of 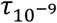 lies before 1981 (Fig. 4B), which suggests that zebra mussels were likely introduced before 1981 and went undetected for at least five years until late 1986 (Carlton 2008). This lag between arrival and first detection is a common feature of invasive species, which arises due to detection failure at low abundances (Crooks 2005). We also find that zebra mussels were likely introduced in Lake Erie (Fig. 4D) and not in Lake St. Claire (Hebert et al. 1989). However, within Lake Erie, the statistical model and occurrence data are consistent with multiple regions as possible locations for origin, a feature that also bears out in synthetic data simulation (Fig. 4C). In fact, there is only one in three chance that mussels were introduced near Kingsville, where they were first detected (Carlton 2008) (Fig. 4D). This uncertainty in location of origin stems from uneven spatial sampling as zebra mussel could have been introduced at a less frequently sampled location and later dispersed and detected at a more frequently sampled location.

Next, we numerically performed elasticity analysis to ask how small perturbations in life-history parameters (***A*** and ***B***) change the arrival time. We find that the arrival time (Eq. [10]) is most sensitive to transition probability from age class 1 to 2 (*S*_1_), hitchhiking rate of individuals in age class 2 (*B*_22_), and fecundity of individuals in age class 2 (*F*_2_). Specifically, a 1% change in these parameters (*S*_1_, *B*_22_, and *F*_2_) will slow the expansion rate by approximately 0.1% (Fig. 5A). Thus, targeted management strategies of disrupting these life-history parameters could potentially slow the spread of zebra mussels. Elasticity analysis also reveals another important and often neglected aspect of species spread—not all age groups contribute equally to species dispersal. The stable age distribution for zebra mussels is skewed toward early life stages, while reproductive value is skewed toward later life stages (Fig S4). Cumulatively, these two trends, with the assumption that the hitchhiking rate of mussels is very low for age class 1 (Keevin et al. 1992), suggest that age class 2 is the main contributor to effective dispersal (Fig. 5B).

**Figure 5:**
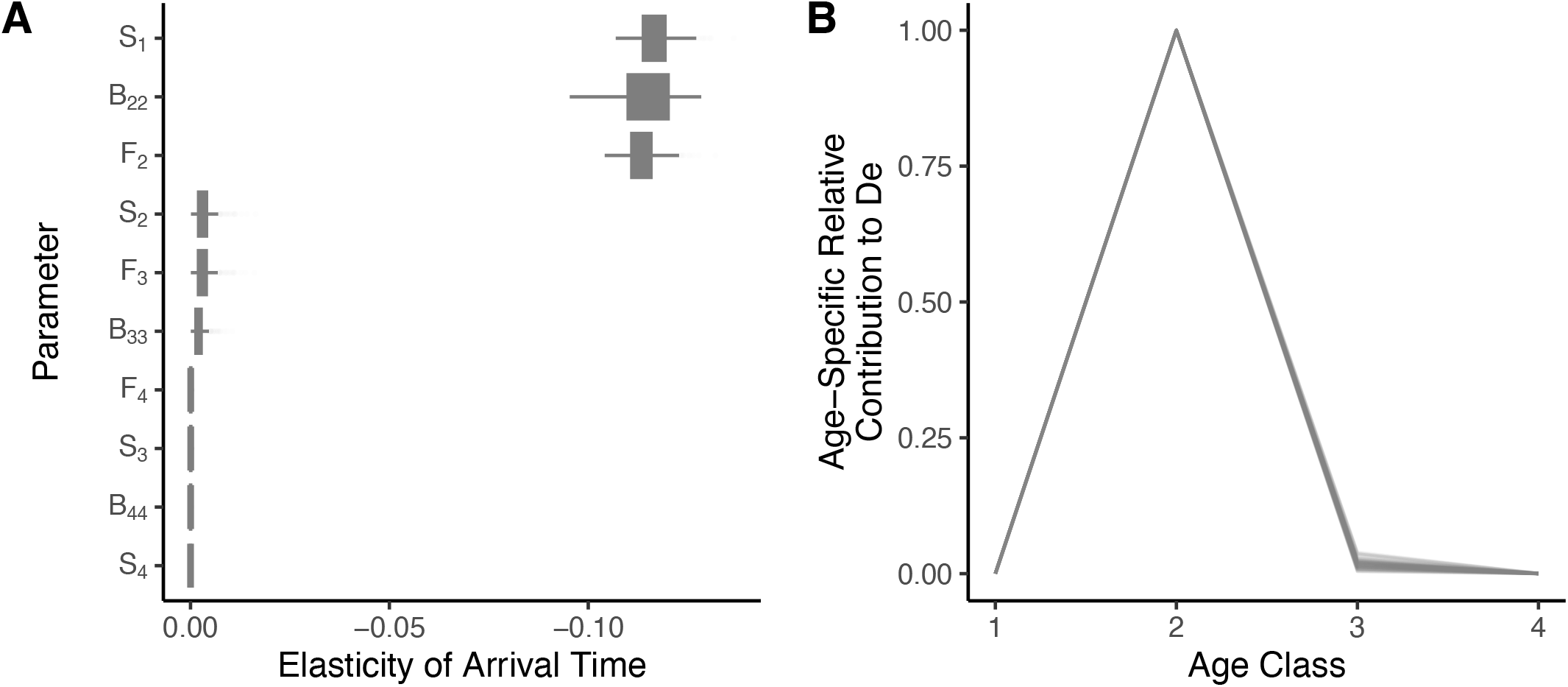
(**A**) Elasticity or proportional sensitivity analysis shows that the arrival time (Eq. [10]) of zebra mussels is most sensitive to transition probability from age class 1 to 2 (*S*_1_), hitchhiking rate of individuals in age class 2 (*B*_22_), and fecundity of individuals in age class 2 (*F*_2_). (**B**) Not all age classes contribute equally to zebra mussel dispersal (Eq. [4]). Age class 2 has the highest relative contribution to effective dispersal, *D*_*e*_.

## DISCUSSION

We show that the metapopulation dynamics of an invader (Eq. [1]) can be simplified into a redistribution matrix with an underlying exponential growth model (Eq. [6]), such that the effective dispersal (*D*_*e*_) and growth rates (*λ*) are a function of life-history parameters (Eqs. [2] and [4]). This hierarchical formulation of the metapopulation model offers several advantages for conceptually understanding drivers of human-mediated invasions.

First, by expressing the dispersal process as a redistribution matrix, we can capture the spatial heterogeneity in the dispersal process and exploit the properties of probability matrices to describe the expansion patterns using the concept of network distances (Eq. [8]). Specifically, the invasion path corresponds, with high probability, to the shortest network path from the origin such that the arrival time is a linear function of the network distance along this path (Eq. [9] and Fig. 1B). Furthermore, visualizing the invasion path on the network space produces a radial wavefront (Fig. 1D), which is geometrically analogous to the radial spread patterns produced by the Gaussian IDE models on the geographical space (Brockmann and Helbing 2013; Kot et al. 1996).

Second, splitting the metapopulation model into spatial and life-history dynamics produces counterintuitive predictions about what life-history tactics make an invader successful. It is widely accepted that species with a high growth rate (Crawley 1986) and propagule pressure (Simberloff 2009) will likely be more successful. However, this assertion ignores that these population-level parameters emerge from life-history traits of individuals that vary in age or ontogenetic stages (Caswell 2001). We show that even when growth rate and propagule pressure are positive, the invasion will fail if the individuals do not disperse at the age when their reproductive value is finite. Corollary, everything else being equal, an invader that disperses at peak reproductive value (e.g., age of maturity) is more likely to overcome demographic stochasticity (Engen et al. 2005) due to higher effective dispersal, *D*_*e*_, and, consequently, expand faster (Eq. [4]). This logic also provides quantitative predictions about how evolution by spatial sorting might proceed at invasion fronts (Shine et al. 2011)—evolution will select for individuals that hitchhike with transportation vessels at the age of peak reproductive value or individuals with higher reproductive value when they hitchhike or both.

Finally, a deeper understanding of the synergy between life-history and spatial dynamics allows one to ask novel ecological questions such as “By how much is it necessary to decrease fecundity at age two to stop an invasion?” or “Is it better to inspect vessels for dispersers at early or later life stages?” (Neubert and Caswell 2000). This knowledge allows managers to identify and target the most responsive life-history parameters to maximize resources (Buhle et al. 2005).

Next, we propose a hierarchical Bayesian model to estimate the parameters of the metapopulation model using commonly available presence-only data. We showed that when dealing with such data, one needs to account explicitly for spatial biases in the data collection procedure (Eqs. [12-14]). Failure to do so can lead to spurious parameter estimates and inferences. For instance, because of uneven spatial sampling, some locations are sampled more often. As a result, it is possible that an invader could have been introduced at a less frequently sampled location and later dispersed and, subsequently, detected for the first time at a location where the sampling was relatively frequent. In such a scenario, using the location of first detection as the location of origin may be inaccurate and produce misleading projections of an invader’s path and arrival time. Our analysis of zebra mussel expansion in North America confirms that this scenario is very likely. As expected, based on early detections of zebra mussels (GBIF.org 2022; USGS 2022) (Fig. S1), the Bayesian model suggests that zebra mussels were likely introduced in Lake Erie (Fig. 4D). However, within Lake Erie, the origin of zebra mussels is uncertain—there is only one in three chance that these mollusks were initially introduced in Kingsville, where they were first detected (Carlton 2008). Even more remarkable is that our statistical model facilitated these inferences with sparse detection data—the survey data only had ∼6% confirmed detections—demonstrating the value of a mechanistic statistical approach to learning from limited and biased datasets.

Our statistical approach also highlights that some parameters cannot be inferred from the occurrence data when multiple combinations of parameters yield the same invasion outcome. We show that these non-identifiable parameters can be resolved by imputing knowledge from previous studies and the dynamics of sister species in the form of priors. We use this technique to constrain zebra mussel’s life-history parameters and carrying capacity, which, in turn, allowed us to make two important inferences. First, the expansion rate of zebra mussels is most sensitive to values of fecundity and dispersal in age class two and survivorship from age one to two (Fig. 5A). Second, zebra mussels were likely introduced in North America before 1981 (Fig. 4B), indicating a minimum lag of five-year between the introduction and first confirmed detection in 1986. Although such lags are a common feature of invasions (Crooks 2005), they are notoriously hard to estimate from data. Thus, using informative priors and explicitly modeling the observation and population processes can allow us to estimate invasion parameters that are otherwise challenging to infer (Hobbs and Hooten 2015).

There are several possible avenues to build on our work. In the metapopulation model, we ignore niche constraints imposed by spatial variation in environmental conditions (Peterson et al. 2011). The environment can also vary in time due to diurnal and seasonal fluctuations (Lande and Orzack 1988). Variation can also stem from demographic stochasticity associated with reproduction and dispersal (Engen et al. 2005). Mechanistically incorporating these sources of variation in population dynamics may provide us with a more realistic understanding of invasion patterns.

On the statistical front, too, many opportunities exist to improve inferences. We showed that one can obtain reliable parameter estimates by explicitly modeling the observation process. However, these improvements come at the expense of precision. To improve precision, we need standardized biodiversity sampling, which currently exists for only a few taxa due to high cost (Pardieck et al. 2020; Pollard et al. 1994). Standardized species observations can be automated with developments in e-DNA, audio and video sensor technology, and advances in computation (Besson et al. 2022; Keitt and Abelson 2021). Such automated schemes are scalable, cost-effective, and can collect standardized species data at broad spatial scales with high temporal resolution. In congruence, to use these data sources in our Bayesian models, we also need to develop observation models that can link these novel datasets to latent parameters of the invasion model.

There is also a need to improve data collection on human transportation pathways for species whose dispersal is tightly associated with specific transportation mechanisms. Examples of such pathways include the spread of insects through the transport of firewood (Solano et al. 2021) and plant material (Fenn‐Moltu et al. 2023), the introduction of exotic animals through international pet trade (Gippet and Bertelsmeier 2021), and the transport of mosquitoes with used tires (Yee 2008). However, when transportation data is absent, we can model the weights of the dispersal network using a mechanistic model of human movement (Simini et al. 2012; Stouffer 1940; Zipf 1946).

Finally, the most useful extension of our statistical framework is hierarchically linking the metapopulation model of invasion with a management model (or policy) (Yemshanov et al. 2009), such as establishing checkpoints to intercept hitchhikers (Zook and Phillips 2012) or chemically eradicating an invader (Fernald and Watson 2013). Because Bayesian models provide a principled way to quantify uncertainty in model parameters (e.g., vital rates and abundances), adding a management model within the Bayesian hierarchy allows managers to propagate this uncertainty while evaluating the cost of implementing different management strategies and then use decision theory to choose the most cost-effective strategy (Dorazio and Johnson 2003; Williams and Hooten 2016).

## Author Contribution

NG conceived and designed the study. AML and NG procured the port call data from Marine Traffic. NG constructed and analyzed the metapopulation and statistical models with inputs from THK, KSK, and MBH. NG wrote the paper with feedback from AML, THK, KSK, CB, and MBH.

## Data and Code Availability

The R and Stan code has been deposited at GitHub (https://github.com/nikunj410/Zebra_Mussels.git).

## Acknowledgment

We thank Upasana Sarraju for creating the illustration (Fig. 2). We also thank Joseph Carver and Randal Holt for collecting metadata on the shipping network from MarineTraffic. NG acknowledges support from the Stengl-Wyer and continuing fellowships from The University of Texas at Austin. Timothy Keitt acknowledges support from the Stengl-Wyer Endowment and the Office of the Vice President for Research, Scholarship, and Creative Endeavors at The University of Texas at Austin and from the National Science Foundation (BCS-2009669). MBH acknowledges support from NSF 2222525 for methodological aspects of this research. KSK was supported through an NIH grant R01-GM138530. CB was supported by the SEFRI-funded project SPREAD (MB22.00086). AML was supported by grant EVA4.0, no. CZ.02.1.01/0.0/0.0/16_019/0000803 financed by OP RDE. MarineTraffic provided the port call data.

## Notes

https://github.com/nikunj410/Zebra_Mussels.git

### Competing Interest Statement

The authors have declared no competing interest.

