## Supplementary material for "A mechanistic statistical approach to infer invasion characteristics of human-dispersed species with complex life cycle": SI

#### Metapopulation model: Formulation

To model zebra mussel expansion, we consider an age-structured metapopulation model with $k$ locations, such that the population dynamics at location $i$ is given by (Hunter and Caswell 2005):

$$\boldsymbol{N}_{t+1,i} = \boldsymbol{A}\left[\boldsymbol{N}_{t,i} + \gamma \boldsymbol{B} \sum_{j \neq i}^{k} q_{ji} \boldsymbol{N}_{t,j} - q_{ij} \boldsymbol{N}_{t,i}\right]. \quad \text{[S1]}$$

Here, $\boldsymbol{N}_{t,i} = \begin{bmatrix} n_{t+1,i}^{1} & n_{t+1,i}^{2} & \cdots & n_{t+1,i}^{a} \end{bmatrix}^{T}$ is a column vector that contains the number of zebra mussels in $a$ age classes at location $i$ and time $t$, $\gamma q_{ij}$ is the number of ships that move from location $i$ to $j$ in one time-step, and

$$\boldsymbol{A} = \begin{pmatrix} F_1 & F_2 & \cdots & F_{a-1} & F_a \\ S_1 & 0 & \cdots & 0 & 0 \\ 0 & S_2 & \ddots & 0 & 0 \\ 0 & 0 & \cdots & S_{a-1} & S_a \end{pmatrix} \quad \text{[S2]}$$

is the Leslie matrix that captures the age-dependent life-history schedules of zebra mussels (Leslie 1945). The elements $F_{i \neq a}$ and $S_{i \neq a}$ are the fecundity and survivorship probability of individuals in age-class $i$, whereas $F_a$ and $S_a$ are the fecundity and survivorship probability of individuals in age class $a$ and above. We capture age-dependent dispersal rate using a diagonal hitchhiking matrix

$$\boldsymbol{B} = \mathrm{diag}([B_{11} \quad B_{22} \quad \cdots \quad B_{aa}]^{T}) \quad \text{[S3]}$$

such that the element $B_{ii}$ corresponding to the hitchhiking rate of individuals in age class $i$.

#### Metapopulation model: Results

To simplify population dynamics, we assume that the dispersal rate is much smaller than the growth rate of the species. More specifically, we assume that immigration and emigration do not substantially change the long-term proportion of individuals in different age classes from the stable-age distribution, $\boldsymbol{w}$. This approximation allows us to express the population at location $i$ as a scalar multiple of $\boldsymbol{w}$, such that $\boldsymbol{N}_{t,i} \approx x_{t,i}\boldsymbol{w}$. Substituting this approximation in Eq. S1 yields

$$x_{t+1,i}\boldsymbol{w} = A\left[x_{t,i}\boldsymbol{w} + \gamma \boldsymbol{B} \sum_{j \neq i}^{k} q_{ji} x_{t,j} \boldsymbol{w} - q_{ij} x_{t,i} \boldsymbol{w}\right]. \quad \text{[S4]}$$

By multiplying Eq. S4 with $\boldsymbol{v}^{T}$ (reproductive value) from the left and then dividing it by $\boldsymbol{v}^{T}\boldsymbol{w}$, we get

$$x_{t+1,i} = \lambda\left[x_{t,i} + D_e \sum_{j \neq i}^{k} q_{ji} x_{t,j} - q_{ij} x_{t,i}\right], \quad \text{[S5]}$$

where $D_e = \gamma \boldsymbol{v}^{T}\boldsymbol{B}\boldsymbol{w}/\boldsymbol{v}^{T}\boldsymbol{w}$ is the effective dispersal. For notational simplicity, we can re-express Eq. S5 as

$$\boldsymbol{x}_{t+1}^{T} = \lambda\, \boldsymbol{x}_{t}^{T} \boldsymbol{P}, \quad \text{[S6]}$$

where $\boldsymbol{x}_{t}^{T} = [x_{t,1} \quad x_{t,2} \quad \cdots \quad x_{t,k}]$ and

$$P_{ij} = \begin{cases} D_e q_{ij}, & i \neq j \\ 1 - D_e \sum_{\xi \neq i} q_{i\xi}, & i = j \end{cases} \quad \text{[S7]}$$

is the $ij^{\text{th}}$ element of the redistribution (probability) matrix $\boldsymbol{P}$ (Kulkarni 2016). Note that the low dispersal rate assumption implies $P_{ij} \ll 1$ when $i \neq j$ and $P_{ij} \approx 1$ when $i = j$.

Solving the recursive relationship in Eq. [S6], we can show that

$$\boldsymbol{x}_t^T = x_{t_0} \lambda^t \, \boldsymbol{e}_n^T \boldsymbol{P}^t, \quad \text{[S8]}$$

where $x_{t_0} = \boldsymbol{v}^T \boldsymbol{N}_{t_0,n} / \boldsymbol{v}^T \boldsymbol{w}$ is the effective initial population and $\boldsymbol{e}_n^T$ is a row vector with all zero elements except at index $n$, where it takes value one. Using Chapman–Kolmogorov identity (Kulkarni 2016), we expand $\boldsymbol{e}_n^T \boldsymbol{P}^t$ to show that the zebra mussel abundance at location $m$ (destination) at time $t$ through all possible paths is

$$x_{t,m} = x_{t_0} \lambda^t \sum_{l_1=1}^{k} \sum_{l_2=1}^{k} \dots \sum_{l_{t-1}=1}^{k} P_{nl_1} P_{l_1 l_2} \dots P_{l_{t-1} m}. \quad \text{[S9]}$$

To simplify Eq. [S9], we define a set of all paths, $\Gamma_{nm}$, from $n$ (origin) to $m$ (destination) in the order of decreasing probability that a zebra mussel takes that path to arrive at $m$. We label these paths as $\Gamma_{nm}^{SP_i}$, where the superscript $SP_i$ denotes the $i^{th}$ path in the ordered set. Since human-transportation networks are highly heterogeneous, as a first approximation, we assume that $\Gamma_{nm}^{SP_1}$ is the invasion path (Brockmann and Helbing 2013; Iannelli et al. 2017). Keeping the terms corresponding to $\Gamma_{nm}^{SP_1}$ in Eq. S9, we get

$$\begin{aligned} x_{t,m} &\approx \binom{t}{n_{SP_1}} x_{t_0} \lambda^t \, D_e^{n_{SP_1}} \wp_{nm}^{SP_1} \\ &\approx \frac{t^{n_{SP_1}}}{n_{SP_1}!} \, x_{t_0} \lambda^t D_e^{n_{SP_1}} \wp_{nm}^{SP_1}, \end{aligned} \quad \text{[S10]}$$

where

$$\wp_{nm}^{SP_1} = \prod_{(i,j) \in \Gamma_{nm}^{SP_1}} q_{ij} \quad \text{[S11]}$$

and $n_{SP_1} (\ll t)$ is the number of steps taken along $\Gamma_{nm}^{SP_1}$. Next, we set the initial population to one (*i.e.*, $x_{t_0} =$ 1) and consider a location is invaded when invader abundance is one. Accordingly, we can show that the arrival time of the species at location $m$ is

$$\begin{aligned} \tau_{nm}^{SP_1} &\approx \frac{n_{sp}}{\log \lambda} \mathbb{W} \left( \frac{\log \lambda}{n_{sp} D_e} \left[ \frac{n_{SP_1}!}{\wp_{nm}^{SP_1}} \right]^{\frac{1}{n_{SP_1}}} \right) \\ &= \frac{D_{nm}^{SP_1}}{\log \lambda}. \end{aligned} \quad \text{[S12]}$$

where $\mathbb{W}$ is the Lambert W function (Gautreau et al. 2008; Lehtonen 2016) and $D_{nm}^{SP_1}$ is the total network distance from location $n$ to $m$ along path $\Gamma_{nm}^{SP_1}$. Using the properties of Lambert W function (Lawley 2020), we can approximate $D_{nm}^{SP_1}$ as

$$\begin{aligned} D_{nm}^{SP_1} &\approx n_{SP_1} \log \frac{\log \lambda}{D_e} - n_{SP_1}\gamma_e - \log \wp_{nm}^{SP_1} \\ &= \sum_{(i,j)\in\Gamma_{nm}^{SP_1}} \log \frac{\log \lambda}{D_e} - \gamma_e - \log q_{ij} \\ &= \sum_{(i,j)\in\Gamma_{nm}^{SP_1}} \mathbb{W}\left(\frac{\log \lambda}{D_e q_{ij}}\right) \\ &= \sum_{(i,j)\in\Gamma_{nm}^{SP_1}} d_{ij} \end{aligned} \qquad [S13]$$

Here, $\gamma_e$ is the Euler-Mascheroni constant and $d_{ij}$ is the network distance between location $i$ and $j$ that share an edge.

Next, to account for alternate invasion paths, we consider the contribution to $x_{t,m}$ by all paths:

$$x_{t,m} \approx \sum_i \lambda^t \frac{t^{n_{SP_i}}}{n_{SP_i}!} D_e^{n_{SP_i}} \wp_{nm}^{SP_i}. \qquad [S14]$$

In general, Eq. [S14] cannot be solved analytically for arrival time. However, we can simplify our analysis by considering paths for which $n_{SP_i} = n_{SP_1}$. This is a reasonable assumption because as $n_{SP_i}$ increases, $\wp_{nm}^{SP_i}/n_{SP_i}!$ decreases drastically. This happens because of the low dispersal assumption—the product in Eq. S11 gets exponentially smaller as $n_{SP_i}$ increases. This allows us to express $x_{t,m}$ as

$$x_{t,m} \approx \lambda^t \frac{t^{n_{SP_1}}}{n_{SP_1}!} D_e^{n_{SP_1}} \sum_i \wp_{nm}^{SP_i}. \qquad [S15]$$

Following the calculations for the shortest path in Eq. [S13], we can show that the most probable network distance (probabilistic average) from $n$ to $m$ is

$$\begin{aligned} D_{nm}^{MP} &= n_{SP_1} \log \frac{\log \lambda}{D_e} - n_{SP_1}\gamma_e - \log \sum_i \wp_{nm}^{SP_i} \\ &= -\log \sum_i e^{-D_{nm}^{SP_i}}. \end{aligned} \qquad [S16]$$

### Supplementary Figures

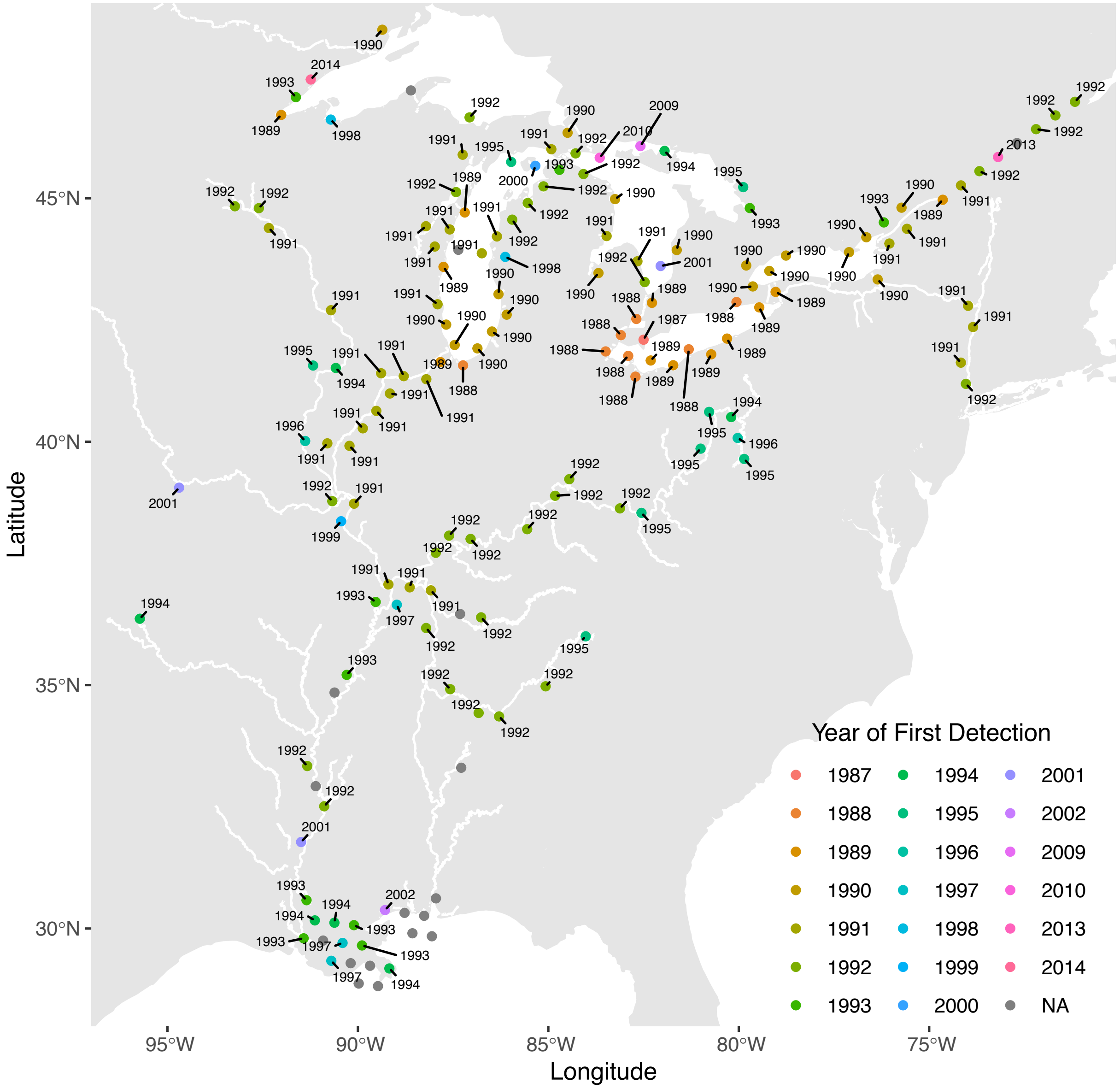


**Figure S1:** First detection records of zebra mussels in North America (GBIF.org 2022; USGS 2022). Data suggests that zebra mussels were first detected near Kingsville (Lake Erie) in 1987 and subsequently colonized inland waterways by 2010. Note that the first detection event in our dataset was in 1987 and not 1986, as stated in the main text. This is because we removed the 1986 data point during quality control. However, the omission of the 1986 data point does not change our results because the zebra mussels were reported in December of 1986, which is only a month away from 1987 when zebra mussels were reported for the second time near Kingsville. Grey points denote locations where zebra mussels were not detected from 1980 to 2019.

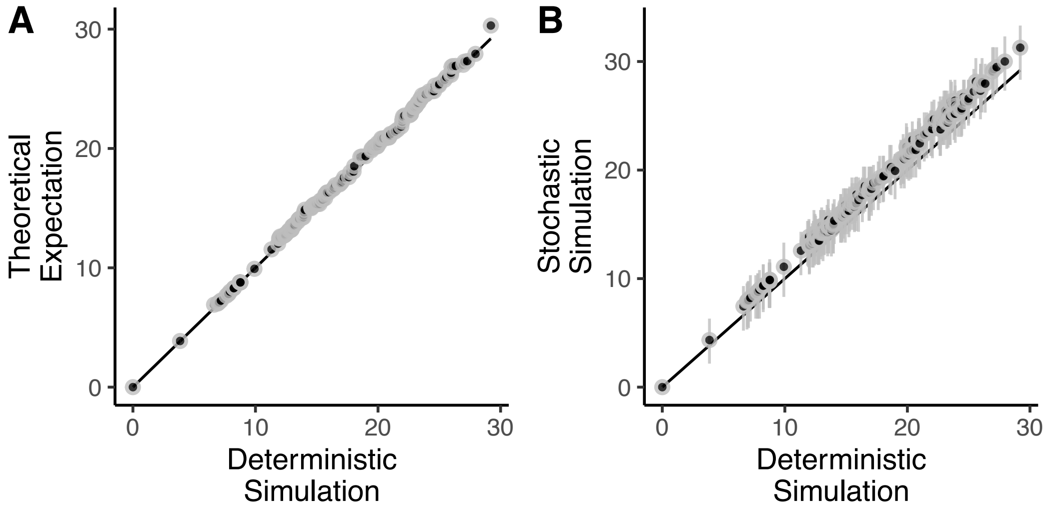


**Figure S2:** The plots compare the species arrival time obtained from deterministic simulation (Fig 1) with theoretically predicted arrival time from Eq. [S12] (**A**) and stochastic simulation in which fecundity, survival, and dispersal are stochastic events (**B**). We find a good match between deterministic simulation, and theoretical results (about 2% error). However, we see that the mean time of species arrival for stochastic simulation is slightly more than deterministic simulation (about 9% error). The back line represents a perfect match.

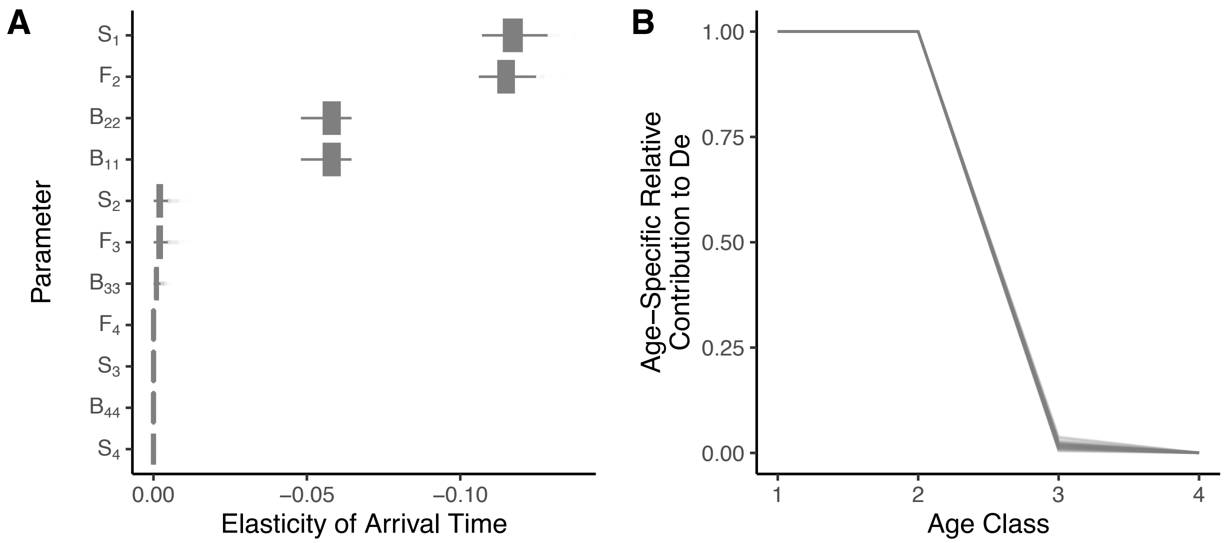


**Figure S3:** The plots show the elasticity analysis and age-dependent contribution to effective dispersal for the alternative hypothesis in which we assume a constant hitchhiking rate. (**A**) As before, elasticity analysis shows that the arrival time of zebra mussels is sensitive to transition probability from age class 1 to 2 ($S_1$), hitchhiking rate of individuals in age class 2 ($B_{22}$), and fecundity of individuals in age class 2 ($F_2$). In addition, we find that the expansion rate is also sensitive to the hitchhiking rate of individuals in age class 1 ($B_{11}$). (**B**) For the alternative hypothesis, both age class 1 and age class 2 have the highest relative contribution to effective dispersal, $D_e$.

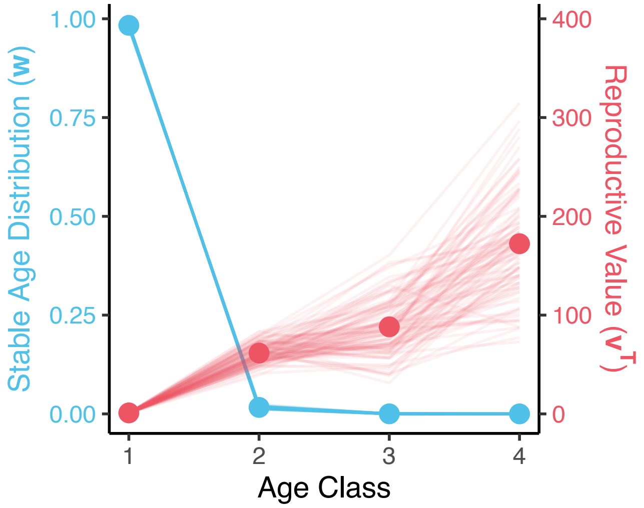


**Figure S4:** Posterior distribution of stable-age distribution ($\boldsymbol{w}$) and reproductive value ($\boldsymbol{v}^{\boldsymbol{T}}$) of zebra mussels. The stable-age distribution is skewed towards early life stages, while reproductive value is skewed towards later life stages. The solid points represent mean values.

### Supplementary Tables

| Sub-model | Sampling statements and relationships | Priors |
|---|---|---|
| Observer model | $y_{t,i} \sim \text{Bernoulli}\left(p_s^i \, p_d^{t,i}\right)$<br>$p_s^i = \text{logit}^{-1}(\beta_i)$<br>$[\beta_1 \; \beta_2 \; \dots \; \beta_k]^T \sim \text{MVNormal}\left(\mu_\beta, \boldsymbol{K}\right)$<br>$K_{ij} = \eta^2 e^{-D_{ij}^2/2\rho^2} + \delta_{ij}\sigma^2$<br>$p_d^{t,i} = 1 - e^{-x_{t,i}\pi_p}$ | $\rho \sim$ Exponential (2)<br>$\sigma \sim$ Exponential (2)<br>$\eta \sim$ Exponential (2)<br>$\beta_i \sim$ Normal (0, 1)<br>$\mu_\beta \sim$ Normal (0, 1.5)<br>$K\pi_p \sim$ Exponential (1) |
| Spatial dynamics model | $\boldsymbol{x}_{\boldsymbol{0}}^T = x_{t_0}\boldsymbol{e}_{\boldsymbol{n}}^T$<br>$\boldsymbol{x}_{\boldsymbol{t+1}}^T = \boldsymbol{x}_{\boldsymbol{t}}^T \boldsymbol{G}_{\boldsymbol{t}} \boldsymbol{P}$<br>$\boldsymbol{G}_{\boldsymbol{t}} = \text{diag}\left(\left[\lambda^{1-\frac{x_{t,1}}{K}} \quad \lambda^{1-\frac{x_{t,2}}{K}} \quad \dots \quad \lambda^{1-\frac{x_{t,k}}{K}}\right]^T\right)$<br>$P_{ij} = \begin{cases} D_e q_{ij}, & i \neq j \\ 1 - D_e \sum_{\xi \neq i} q_{i\xi}, & i = j \end{cases}$ | $n \sim$ DiscreteUniform $(1, k)$<br>$\log\left(x_{t_0}/K\right) \sim$ Normal (-5, 2.5)<br>$\log_{10}(K) \sim$ Uniform (13.4, 14.4)<br>$D_e/3 \sim$ Exponential (1) |
| Life-history model | $\boldsymbol{A} = \begin{pmatrix} 0 & \sigma_0 f_1 & \sigma_0 f_2 & \sigma_0 f_3 \\ \sigma_1 & 0 & 0 & 0 \\ 0 & \sigma_2 & 0 & 0 \\ 0 & 0 & \sigma_3 & \sigma_4 \end{pmatrix}$<br>$\boldsymbol{A}\boldsymbol{w} = \lambda \boldsymbol{w}$<br>$\boldsymbol{v}^T\boldsymbol{A} = \lambda \boldsymbol{v}^T$<br>$D_e = \Upsilon(\boldsymbol{v}^T\boldsymbol{B}\boldsymbol{w}/\boldsymbol{v}^T\boldsymbol{w})$<br>$\boldsymbol{B} = \text{diag}([0 \quad B \quad B \quad B]^T)$ | $10^{-2}\sigma_0 \sim$ Normal (1.50, 0.25)<br>$10^{-1}\sigma_1 \sim$ Normal (6.50, 1.80)<br>$10^{-1}\sigma_2 \sim$ Normal (5.00, 1.50)<br>$10^{-1}\sigma_3 \sim$ Normal (2.50, 0.75)<br>$10^{-2}\sigma_4 \sim$ Normal (8.80, 0.62)<br>$10^{5}f_1 \sim$ Normal (1.50, 0.50)<br>$10^{5}f_2 \sim$ Normal (2.40, 0.81)<br>$10^{5}f_3 \sim$ Normal (5.00, 1.80) |

**Table 1:** The table provides the list of sampling statements, relationship between parameters, and priors. Based on published literature, we constructed informative priors for life-history parameters (Casagrandi et al. 2007) and carrying capacity (Cross et al. 2011). For all other parameters, we used weakly informative priors. For $\boldsymbol{B}$, we considered an alternate hypothesis—$\boldsymbol{B} = \text{diag}([B \quad B \quad B \quad B]^T)$ (see Fig. S3).

USGS. 2022. *Dreissena polymorpha* (Pallas, 1771): U.S. Geological Survey, Nonindigenous Aquatic Species Database. Gainesville, FL, United States Geological Survey.
